# Let-7i-5p regulation of cell morphology and migration through distinct signaling pathways in normal and pathogenic fibroblasts from urethra

**DOI:** 10.1101/330332

**Authors:** Kaile Zhang, Jun Chen, Shukui Zhou, Ranxin Yang, Ying Wang, Qiang Fu, Rong Chen, Xiaolan Fang

## Abstract

Pelvic fracture urethral distraction defects (PFUDD) is a common disease that could severely affect patients’ life quality, yet little is known about the molecular mechanism associated with pathogenic fibrosis in PFUDD. In this study, we found that let-7i-5p could regulate different cellular events in normal and pathogenic fibroblasts through three distinct signaling pathways. Interestingly, those regulations are compromised during the translation from mRNA to protein, and partially based on pathogenic status of the fibroblasts. By analyzing the molecular mechanism associated with its function, we conclude that let-7i-5p plays an essential role in regulating cell shape and tissue elasticity, cell migration, cell morphology and cytoskeleton, and could serve as a potential target for clinical treatment of urethral stricture patients.

## Introduction

Pelvic fracture urethral distraction defects (PFUDD) is a common disease that could severely affects patients’ life quality, largely due to excessive fibrosis and associated urethral stricture[1]. Fibrosis is a key factor responsible for pathologic changes related to urethral stricture (in both primary or recurrent diseases)[2, 3]. Over the last few decades, microRNAs and their regulation of fibrosis have been studied in many specific organs, such as liver, heart, skin, kidney and lung[4-11]. The major interests are in miR-29 and TGF-beta signaling pathway, focusing on their role of molecular regulation of fibrosis and/or associated excessive extracellular matrix deposition[4, 9]. miRNAs in specific diseases, such as idiopathic pulmonary fibrosis (IPF), together with their functions in epithelial-mesenchymal transition (EMT) and trans-differentiation, have also been studied [12]. Thus, it would be extremely helpful to further understand molecular mechanisms and related miRNA signaling involved in PFUDD-associated fibrosis, in order to discover novel targets to prevent PFUDD by suppressing urethral stricture.

Our group recently performed molecular profiling of microRNAs in PFUDD patients and summarized a few candidates that may serve regulatory functions in fibrosis[1]. We found that miR-29 expression is medium in normal and PFUDD patients’ scar tissues, and miRNA sequencing data showed that expression of hsa-miR-29b-3p and hsa-miR-29c-3p were both slightly downregulated in scar tissues from PFUDD patients [1] (although the differences are significant, as the fold is 0.64 and 0.69, scar/normal). In addition, we found that let-7i-5p showed very high expression among all the microRNAs sequenced by microRNA sequencing, and its expression showed an increase in scar tissue comparing to normal tissue (although the difference is not significant). We extended our work of microRNA analysis in PFUDD and discovered let-7i-5p as a novel regulator in different cellular events in normal and fibrotic urethral tissues. We hypothesize that let-7i-5p may serve as an active regulator in cellular events, given its impressive abundance in normal and pathogenic fibroblasts. In this study, we analyzed let-7i-5p by up- or down-regulating its expression in normal human fibroblasts and pathogenic tissues. By evaluating potential molecular targets (COL1A1, COL3A1, ELN, MMP1, VIM, FN1, ACTIN, TGFBR1 and TIMP1) at mRNA and protein levels, we testified the regulation by let-7i-5p in cell shape and tissue elasticity, cell migration, cell morphology and cytoskeleton through several signaling pathways.

## Material and Methods

### Urethral scar samples

The study was approved by the Ethics Committee of Shanghai Sixth People’s Hospital. Consents were obtained from all of the patients to use their samples in scientific research. Scar tissues in urethra (Human scar fibroblasts, or HSF) were harvested from PFUDD patients undergoing urethroplasty (n=5). All five subjects are males (gender ratio is 100% male) with ages ranging from 16 to 59. Patients’ baseline information was summarized in Table I. The etiology of patients with urethral stricture was PFUDD. All the participating patients underwent primary surgery. The mean length of stricture is 1.5 cm and the locations were all at the membranous segment of urethra. Samples were harvested after surgery, sectioned and stored at −80 °C until the process of RNA extraction.

**Table I.**
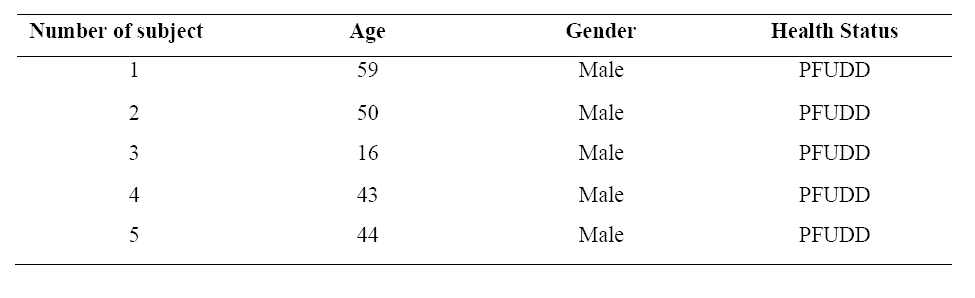
Patient baseline characteristics

### Human cell line

Normal human foreskin fibroblasts (HFF, Catalog# SCSP-106) were provided by Stem Cell Bank, Chinese Academy of Sciences.

### Cell transfection

HSF and HFF cells were growing in 10 cm dishes in Dulbecco’s MEM (DMEM, Gibco, Cat.# 12100-046, Carlsbad, CA, USA) supplemented with Fetal Bovine Serum (FBS) (Gibco, Cat.#10099-141, Carlsbad, CA, USA) to 10% by volume and Penicillin-Streptomycin (100 U/ml) (Gibco, Cat.# 15140122, Carlsbad, CA, USA). 80-90% confluency cells were then detached using Trypsin (Gibco, Cat.# 25200056) and plated at 1 ×; 10^5^ cells/ml in 6-well dishes (2ml/well), and incubate at 37°C overnight. The cells were transfected with lentiviral constructs (empty control construct, customized lenti-KD miRNA and lenti-OE miRNA from GENECHEM, Shanghai, China) to overexpress or knock down of let-7i-5p according to the manufacturer’s protocol. Transfection mixture was replaced by 2ml DMEM (10% FBS) medium after 12 hours. Transfected cells were grown for 72 hours before imaging.

### Imaging

To confirm stable expression of each transfected construct, GFP expression of transfected cells was observed and evaluated by an Olympus IX70 microscope under fluorescent channel. For regular bright field imaging (for Transwell assay), samples were imaged with the Olympus IX70 microscope under bright light channel.

### Cell migration assay

Inserts with 8 μM pore size (Corning-Costar, Lowell MA) were used with matching 24-well transwell chambers. Cells were suspended in serum-free DMEM medium and adjusted to 2 ×; 10^5^ cells/ml. 100-150µl cell suspension were placed in the upper chambers. The lower chambers were filled with DMEM medium with 10% FBS (600-800 µl /well). Cells were incubated at 37C for 24 hours, the inserts were removed and inner side was wiped with cotton swaps. The inserts were then fixed in methanol for 30 minutes at room temperature, and stained with crystal violet solution (Cat.#A100528-0025, Sangon Biotech Shanghai, China) for 15-30 minutes and were peeled off after washing and mounted on the slides. The migrated cells were imaged with a OLYMPUS IX70 microscope using bright light channel. Six cell types (HFF (NC (negative control)/SI ((for siRNA-led knockdown)/OE (overexpression), or FNC/FSI/FOE and HSF (NC (negative control)/SI ((for siRNA-led knockdown)/OE (overexpression), or SNC/SSI/SOE) were analyzed and triplicate experiments were done for all the cell types.

### Reverse transcription (RT)-qPCR for miRNAs and targeted genes

Total RNA was extracted following standard protocol by Servicebio, Inc. (Wuhan Servicebio Technology Co., Ltd, Wuhan, Hubei, China). The primers used for PCR were designed with Primer Premier software (version 5.0; Premier Biosoft International, Palo Alto, CA, USA) (primer sequence details are summarized in Table II). cDNA synthesis was performed on a GeneAmp PCR System 9700 (Applied Biosystems; Thermo Fisher Scientific, Inc., Waltham, MA, USA) following the manufacturer’s instructions (RevertAid First Strand cDNA Synthesis kit, Cat.# K1622, Thermo Fisher Scientific, Inc., Waltham, MA, USA). qPCR was performed on a ViiA 7 Real-time PCR System (Applied Biosystems; Thermo Fisher Scientific, Inc.) using a PowerUp SYBR Green Master Mix (Cat.# A25778, Thermo Fisher Scientific, Inc.). The PCR thermal procedure is 1) 95°C, 10min, 2) 95°C, 15s->60°C, 60 s, 40 cycles. The fold change for each miRNA was calculated using the 2^−ΔΔCq^method [13]. U6 expression level was used to normalize the mRNA expression data. Expression in six cell types (HFF (NC/SI/OE) and HSF (NC/SI/OE) were analyzed and triplicate experiments were done for all the cell types.

**Table II.**
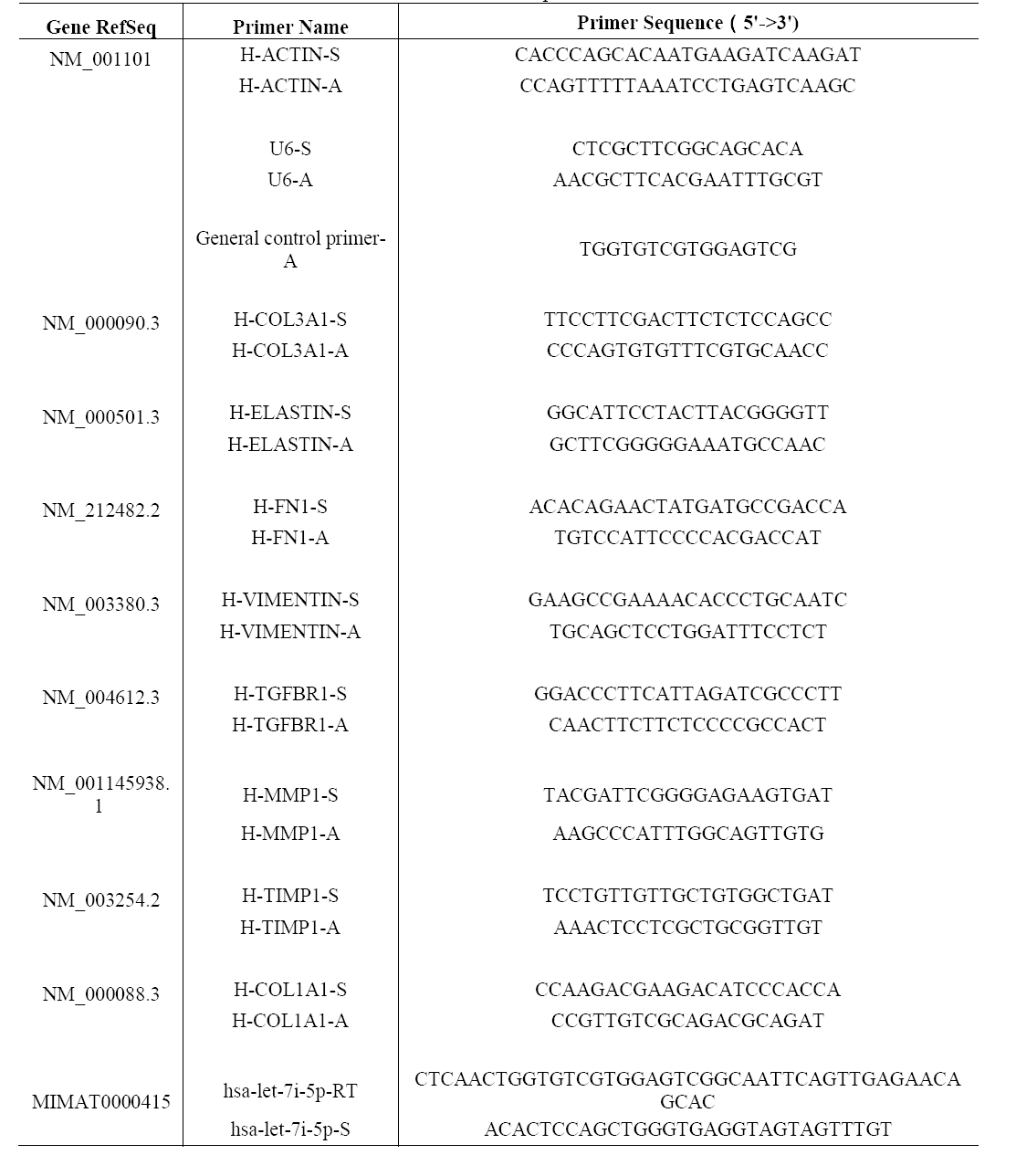
RT-PCR primers

### Western Blotting

Whole protein lysate was extracted using RIPA Lysis buffer (Cat.#20101ES60, Yeasen, Shanghai, China) following instructions by the manufacturer. Equal concentration of protein was loaded on 5-10% SDS-PAGE gels and transferred onto a PVDF membrane (Cat.# IPVH000010, MilliporeSigma, Burlington, MA, USA). Primary antibodies for Col1a1, Col3a1, Elastin, Vimentin, MMP1, Fibronectin, TIMP1, Actin, TGFBR1 and GAPDH were listed in Table III. Bands were visualized using horse-radish peroxidase (HRP) conjugated secondary antibodies (Table III) in conjunction with Immobilon ECL Ultra Western HRP Substrate (Cat.#WBKLS0100, MilliporeSigma, Burlington, MA, USA) via ImageQuant LAS 4000mini (GE Healthcare Life Sciences, Pittsburgh, PA, USA). Expression in six cell types (HFF (NC/SI/OE) and HSF (NC/SI/OE)) were analyzed and triplicate experiments were done for all the cell types. AlphaEaseFC software (Genetic Technologies Inc., Miami, FL, USA) was used to analyze the density of electrophoretic Western blot bands by Servicebio, Inc. (Wuhan Servicebio Technology Co., Ltd, Wuhan, Hubei, China). GAPDH expression level was used to normalized the protein expression data. Intensity analysis was done for one of the experiments.

**Table III.**
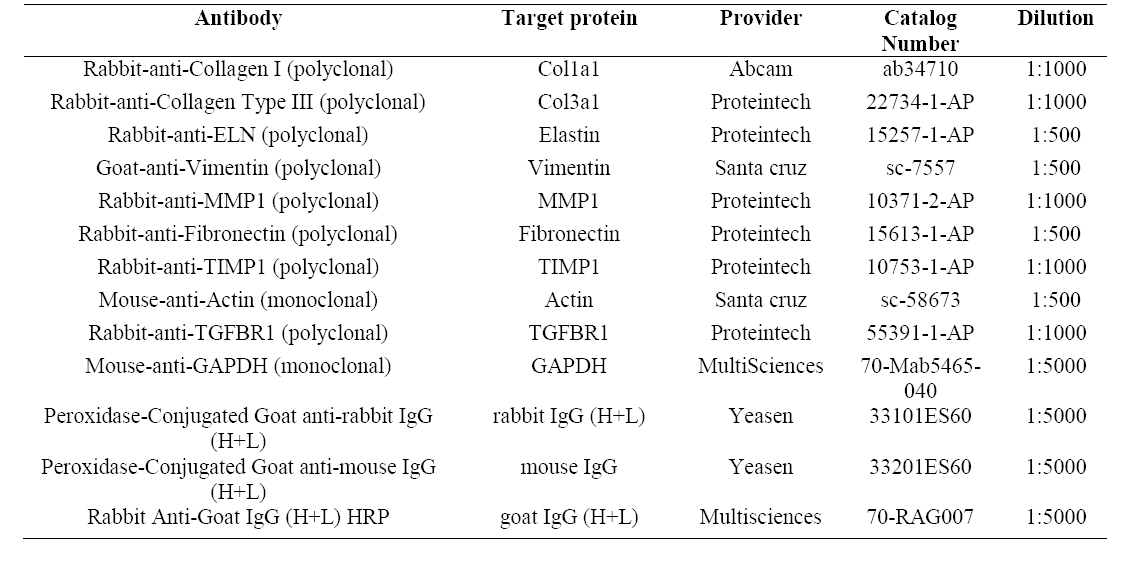
Commercial antibodies

### Enzyme Linked Immunoabsorbent Assay (ELISA)

Supernatant of cell lysates was collected for each of six cell types (HFF (NC/SI/OE) and HSF (NC/SI/OE)), and the levels of MMP2, TGFβ1 and TIMP1 were quantified using ELISA kits as per manufacturers’ instructions (Human Matrix Metalloproteinase 2/Gelatinase A (MMP-2) ELISA Kit, Cat.#CSB-E04675h, CUSABio, Wuhan, Hubei, China; Human TGF-beta1 ELISA Kit, Cat.#EK1811, MultiSciences, Hangzhou, Zhejiang, China; Human TIMP1 ELISA Kit, Cat.#EK11382, MultiSciences).

### Statistical analysis

Student t-test were performed for let-7i-5p expression comparison (unpaired, two tails, heteroscedastic) in six cell types (FNC, FSI, FOE, SNC, SSI and SOE). Student t-test were performed for the standard deviation comparison (paired, two tails) of mRNA and protein expression of COL1A1, COL3A1, ELN, MMP1, VIM, FN1, ACTIN, TGFBR1 and TIMP1.

## Results

### Let-7i-5p regulates cell morphology and motility in normal and pathogenic fibroblasts

Let-7i-5p is a member of Lethal-7 (let-7) microRNA family, which is widely observed and highly conservative across species, from reptiles to mammals (Figure 1A). Let-7 family members were among the first discovered microRNAs and were shown to be an essential regulator of development in *C. elegans*[14], and let-7 microRNA family has been reported to regulate allergic inflammation through T cells[15]. In human tissues, hsa-let-7i-5p showed extremely high expression in thyroid,, and relatively high expression in spinal cord, brain, muscle and vein, with tissue specific index score at 0.905 (indicating a high tissue expression specificity in thyroid) (Figure 1B) [16], suggesting a potential role of hsa-let-7i-5p in metabolism. However, little is known about the role of let-7i-5p or the related molecular mechanism in normal fibroblast or fibrosis-related scar formation.

**Figure 1.**
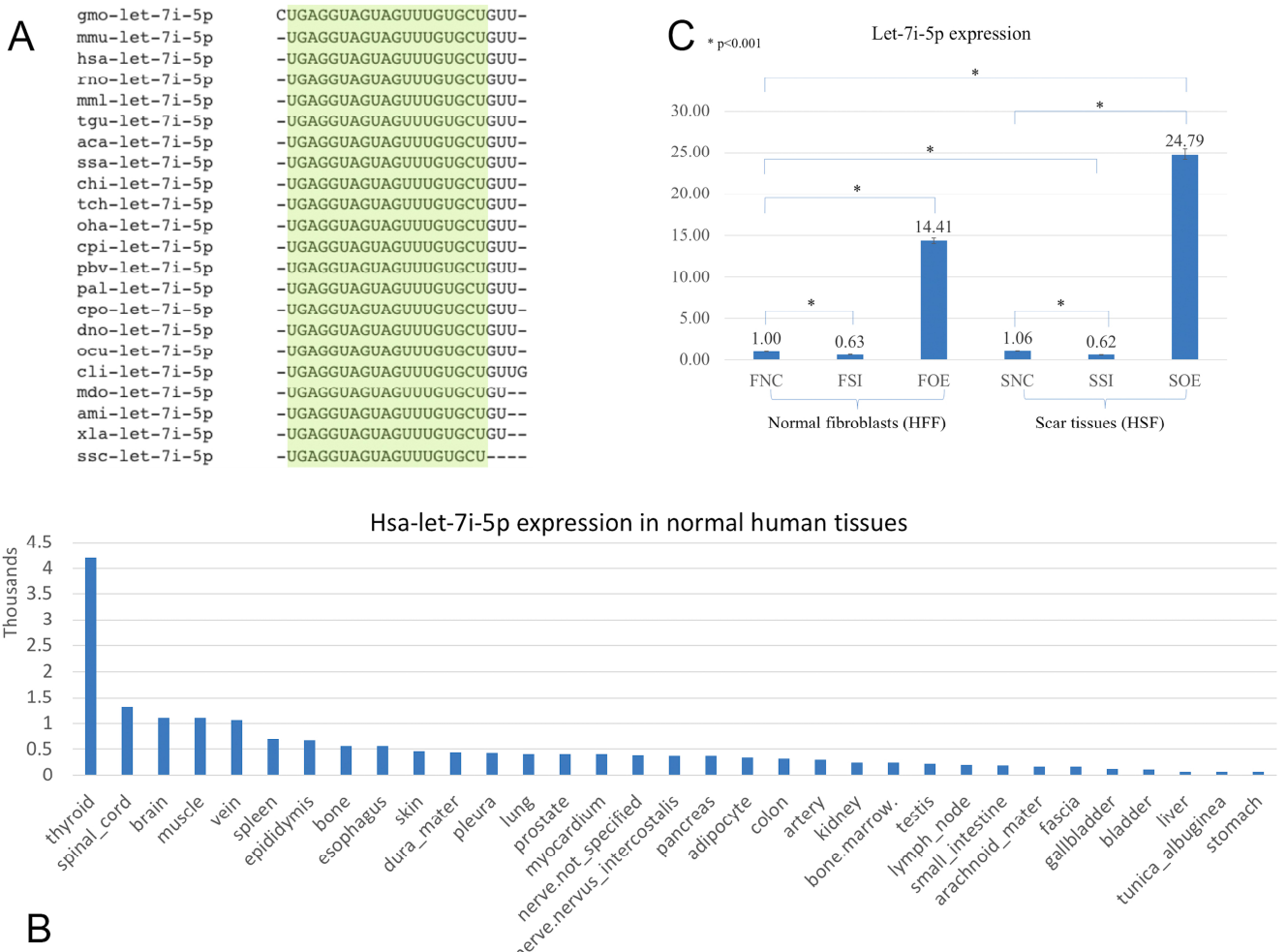
Let-7i-5p is conservative among different species and hsa-let-7i-5p is expressed differentially in normal human tissues. By lenti-viral infection the let-7i-5p expression was manipulated either up or down in normal and scar tissues. A, let-7i-5p sequence comparison across different species. The conservative sequences are highlighted. aca, Anolis carolinensis; ami, Alligator mississippiensis; chi, Capra hircus; cli, Columba livia; cpi, Chrysemys picta; cpo, Cavia porcellus; dno, Dasypus novemcinctus; gmo, Gadus morhua; hsa, Homo sapiens; mdo, Monodelphis domestica; mml, Macaca mulatta; mmu, Mus musculus; ocu, Oryctolagus cuniculus; oha, Ophiophagus hannah; pal, Pteropus alecto; pbv, Python bivittatus; rno, Rattus Norvegicus; ssa, Salmo Salar; ssc, Sus scrofa; tch, Tupaia chinensis; tgu, Taeniopygia guttata; xla, Xenopus laevis. B, hsa-let-7i-5p expression levels in normal human tissues. Data based on two individuals’ microRNA sequencing results (Ludwig et al, *Nucleic Acids Research*, 2016 **44:**3865-3877) and average of normalized value by quantile normalization were used. C, let-7i-5p level was up- and down-regulated in normal and pathogenic fibroblasts by Lenti-viral transfection. *, p<0.001. F, normal fibroblasts (HFF). S, scar tissues. NC, non-transfected control. SI, transfected by lenti-KD miRNA to knock down hsa-let-7i-5p expression. OE, transfected by lenti-OE miRNA to overexpress hsa-let-7i-5p.

Based on a recent miRNA profiling using PFUDD patients’ tissues, we found that let-7i-5p expression was really high in both normal and pathogenic fibroblasts based on miRNA sequencing. To manipulate the knock-down or overexpression of let-7i-5p, the miRNA level was up- and down-regulated in normal (HFF) and pathogenic (HSF) fibroblasts using Lenti-viral transfection and significant expression changes were observed (Figure 1C). Dysregulation of let-7i-5p in normal fibroblasts caused cell morphology changes, yet had little influence on that of pathogenic fibroblasts (Figure 2). Surprisingly, either overexpression or knockdown of let-7i-5p resulted in rounder but more spiky cells. Similar phenotypes were reported to be caused by null-functional Dematin (an actin binding/bundling protein), and was associated with null effect in mutant fibroblasts and impaired wound healing process[17].

**Figure 2.**
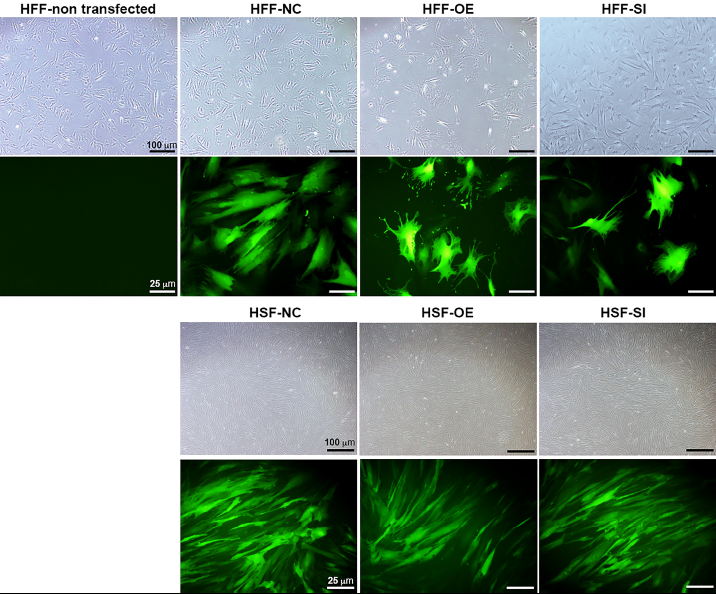
Let-7i-5p level change results in cell morphology changes in normal fibroblasts, but not in pathogenic fibroblasts. Scale bar in bright field images, 100 μm. Scale bar in fluorescent images, 25 μm.

To see whether let-7i-5p could regulate cell motility, we performed cell migration assay for normal and pathogenic fibroblasts. Inhibition of let-7i-5p led to a clear promotion of cell motility, while overexpression of let-7i-5p displayed a severely suppression (Figure 3). The regulation patterns are similar in both normal and pathogenic fibroblasts (Figure 3).

**Figure 3.**
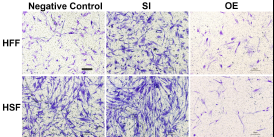
Overexpression of let-7i-5p could suppress migration, while inhibition could promote cell migration in both normal and pathogenic fibroblasts. Scale bar, 50 μm.

### Let-7i-5p exerts distinct regulatory functions in different signaling pathways

We then performed real time quantitative PCR to evaluate the mRNA expression level of potential targets of let-7i-5p to elucidate its function in normal and pathogenic cells. Surprisingly, we found that let-7i-5p regulates signaling molecules through three different patterns (Figure 4A, Table IV).

**Figure 4.**
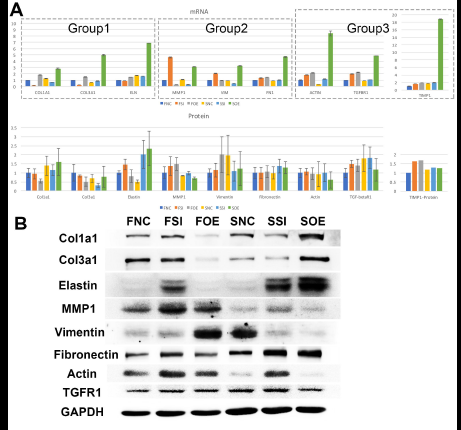
Let-7i-5p regulates signaling molecules in normal and pathogenic fibroblasts. A, quantitative analysis of mRNA and protein expression levels of let-7i-5p’s downstream targets. Group 1, target genes positively regulated by hsa-let-7i-5p in normal fibroblast cells at mRNA level. Group 2, target genes negatively regulated by hsa-let-7i-5p in normal fibroblast cells at mRNA level. Group 3, target genes constitutively upregulated in normal and pathogenic fibroblasts at mRNA level with either suppressed or increased hsa-let-7i-5p expression. Error bar, standard error. mRNA level expression was evaluated by quantitative real-time PCR. Protein level expression was evaluated by western blotting (Col1a1, Col3a1, Elastin, MMP1, Vimentin, Fibronectin, Actin, TGF-betaR1) and ELISA (TIMP1). B, representative western blotting image of protein expressions of target genes.

**Table IV.**
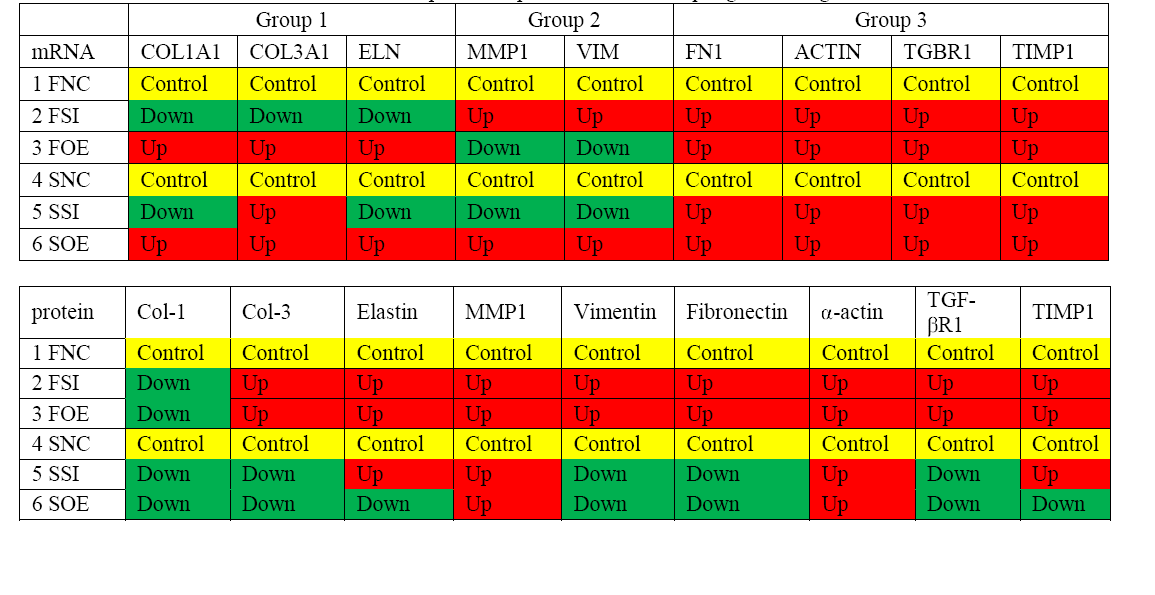
mRNA and protein expressions of let-7i-5p regulated targets

In the first group, at mRNA level, let-7i-5p knockdown resulted in suppression of gene expression of COL1A1, COL3A1 and ELN in both normal and pathogenic fibroblasts, while overexpression of let-7i-5p resulted in significantly increased expression of those genes (Figure 4A, Group 1). This suggests a positive correlation between let-7i-5p and those three genes. Pathogenic status (whether the cell is normal or pathogenic fibroblasts) doesn’t seem to affect this regulation, with the only exception of COL3A1 in let-7i-5p knockdown pathogenic fibroblasts, which had a slight increased expression instead of down regulation.

Interestingly, in the second group, we observed quite opposite regulatory effects on mRNA expression of MMP1 and VIM by let-7i-5p (Figure 4A, Group 2). Knockdown of let-7i-5p significantly enhanced MMP1 and VIM expression, while overexpression of let-7i-5p caused a decrease in expression. Strikingly, this negative regulation is completely reversed in pathogenic fibroblasts, as downregulation of let-7i-5p decreased the expression of MMP1 and VIM, and upregulation of let-7i-5p increased their expression. This strongly suggests that regulation of MMP1 and VIM by let-7i-5p is dependent on the pathogenic status of the fibroblasts.

The third group of regulated genes contains FN1, ACTIN, TGFBR1 and TIMP1, which had increased expression with either knockdown or overexpression of let-7i-5p in both normal and pathogenic fibroblasts (Figure 4A, Group 3). This suggests dysregulated let-7i-5p could boost up the expression of those target genes, regardless of the actual expression level of let-7i-5p (whether it is up- or down-regulated). It is worth noting that the enhanced expression was impressively high for all four target genes in fibrotic cells when let-7i-5p is over-expressed (Figure 4A), suggesting that pathogenic fibroblasts could further amplify the regulatory effect of those genes resulted from let-7i-5p overexpression, while normal cells still maintained a retraining ability to suppress the dysregulation of target genes caused by let-7i-5p level changes.

### Regulatory effects by let-7i-5p were compromised during translation

To confirm whether the regulatory effects at mRNA level were effectively passed on to protein level, we evaluated the protein expression of signaling molecules in normal and pathogenic fibroblasts with manipulated let-7i-5p expression (Figure 4). Quantification of protein expression levels indicated that the effect of let-7i-5p regulation is largely compromised during the translation in normal and pathogenic cells, suggesting an overall neutralization effect (Figure 4A). The huge fold change in mRNA expression (0.18-18.77) shrunk down to a relatively small range for protein levels (0.30-2.34) (Figure 4A). To analyze the variation scale of expression, we compared the standard deviation of each mRNA or protein among all six cell types and significant difference was observed (Figure 5). This suggests that the regulation by let-7i-5p was attenuated during the translation process and resulted in more compromised effect on targeted proteins.

**Figure 5.**
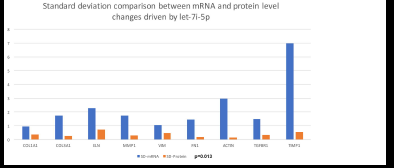
The effect of let-7i-5p regulation is compromised during the translation from mRNA to protein expression in normal cells, suggesting a stabilization effect by translation regulation. Standard deviation is based on the variation of expression level (of each specific mRNA/protein) in all 6 cell groups. p= 0.013 based on student t-test.

At protein level, the regulation of Col1a1, Col3a1 and Elastin is partially retained in HSF, but not in HFF, suggesting that pathogenic fibroblasts may lose part of the ability to neutralize miRNA’s regulatory effect of intracellular targets and impair the cellular homeostasis (Figure 4A, Table IV). The regulation of MMP1 and Vimentin protein levels showed a slightly different pattern comparing to that of mRNA (Figure 4A). In pathogenic fibroblasts, MMP1 expression was reversed from positively regulated at mRNA level to negatively regulated at protein level. MMP2 expression displayed a similar negative pattern at protein level, although the expression level of MMP2 was almost doubled in control pathogenic fibroblasts (SNC) (comparing to normal control cells (FNC)) (Figure 6). Vimentin had an increased expression in normal cells with overexpressed let-7i-5p and pathogenic fibroblasts but remain unchanged in the rest four cell types (Figure 4A). Fibronectin and actin protein levels were comparably unaffected regardless of let-7i-5p regulation (Figure 4A). TGF-beta receptor1(TGFbetaR1) protein level was negatively regulated in normal cells with dysregulated let-7i-5p (knocked down or overexpressing), and in pathogenic fibroblasts it increased in cells with either no change or knock-down of let-7i-5p (Figure 4A). Interestingly, it was suppressed in pathogenic fibroblasts with let-7i-5p overexpression (Figure 4A). This is consistent with the pattern observed in migration assay and may indicate a role of TGFbetaR1 in cell motility regulation. We also evaluated its ligand, TGF-beta1 in corresponding cell types, and observed an opposite pattern of regulation (Figure 6), which indicates the negative feedback regulation of TGF-beta in response to the TGFbetaR1 protein level changes in all six cell types.

**Figure 6.**
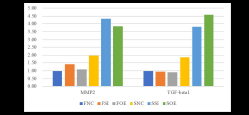
Let-7i-5p regulates MMP2 and TGF-beta1 proteins in normal and pathogenic fibroblasts (ELISA). Protein level expression was evaluated by ELISA.

Overall, the regulation by let-7i-5p is compromised during the translation from mRNA to protein, suggesting a stabilization effect during this process.

## Discussion

### Let-7i-5p serves unique functions in multiple signaling pathways in normal fibroblasts

In normal human fibroblasts, manipulated let-7i-5p expression resulted in positive regulations of COL1A1, COL3A1 and ELN), negative regulations of MMP1 and VIM, and constitutively increased expression of FN1, ACTIN, TGFBR1 and TIMP1 at mRNA level. Given the fact that those molecules are involved in different signaling transduction pathways and corresponding cellular events, our data strongly suggest a multi-functional role of let-7i-5p in normal cells, which could regulate fibroblast proliferation, metabolism, homeostasis and posttranslational modifications, and result in changes of tissue elasticity, cell migration, cytoskeleton and cell morphology (Figure 7).

**Figure 7.**
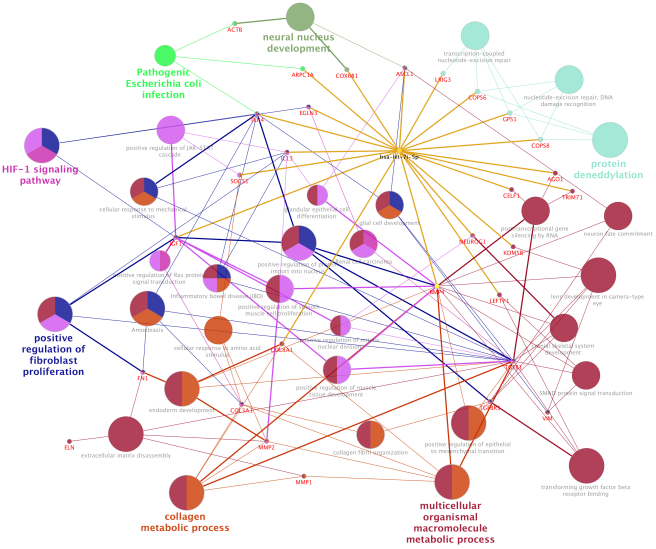
Let-7i-5p regulation mechanism is summarized in distinct functional pathways

### Let-7i-5p regulates collagen metabolic process and tissue elasticity through COL1A1, COL3A1 and possibly COL8A1

The positive regulation of COL1A1, COL3A1 and ELN mRNA expressions in normal fibroblasts was largely retained in pathogenic cells, except for COL3A1, which has a slightly increased expression with decreased let-7i-5p. This indicates that the regulation of those target genes by let-7i-5p was not interrupted in pathogenic fibroblasts, or the functions of let-7i-5p in this specific signaling pathway is independent of the fibrotic status of cells. Since those proteins function together to strengthen and support connective tissues in the body, our data suggest an independent role of let-7i-5p in regulating collagen metabolism and tissue elasticity. This is in concordance with several studies elucidating the association between tissue elasticity and miRNA regulation. For example, in primitive neuroectodermal tumor (PNET) stem cells, tissue elasticity was suggested to promote miRNA silencing and downregulation of target genes [18]. In mouse models, miR-29-3p could suppress the mRNA expression of COL1A1 and COL1A3 either with or without the induction by TGF-β1 and prevent *S. japonicum*-induced liver fibrosis[19]. Also, Col1a1 and Col3a1 were overexpressed during active inflammation and murine colitis induced by 2,4,6-trinitrobenzene sulfonic acid (TNBS) hapten [20]. Given that COL8A1 is a potential target of let-7i-5p, our data suggest a positive regulation pattern of collagen metabolic process at mRNA level by let-7i-5p.

However, at protein level, the regulatory patterns are quite different, with down-regulated expression of Col1a1 and Col3a1 in normal cells. In pathogenic cells, a partial positive regulatory pattern was observed at protein level when let-7i-5p was knocked down, and not much difference was observed with let-7i-5p overexpression. The protein expression changes in normal fibroblasts are more sensitive to let-7i-5p overexpression, while that in pathogenic fibroblasts are more sensitive to let-7i-5p knock-down, reflected by Col1a1 and Col3a1 expression levels (Figure 4A). The regulation of Elastin was reversed to negative pattern in normal fibroblasts at protein level and was constitutively overexpressed in pathogenic cells regardless of whether let-7i-5p was up- or down-regulated (Figure 4A). More evidence is required to determine how the regulation is affected during translation of Elastin protein.

### Let-7i-5p regulates extracellular matrix (ECM) and cell migration through MMP1, MMP2 and Vimentin

Let-7i-5p negatively regulated VIM and MMP1 in normal cells, and this regulation was interrupted in pathogenic fibroblasts, as their mRNA expressions were completely reversed from negative (in normal cells) to positive (in pathogenic cells) regulatory pattern (Figure 4A). This observation indicates that let-7i-5p functions as an upstream regulator of Vim and MMP1 and its regulation is dependent on pathogenic status of the cells.

TGF-β1 and MMP2 protein levels were up-regulated, together with increased cell migratory capability in pathogenic fibroblasts comparing to normal cells (Figure 6). This is consistent with what was reported in human hepatic stellate cells, in which stimulation with TGF-β1 resulted in an increase in migratory capacity and up-regulated MMP-2 activity[21]. However, in pathogenic fibroblasts with overexpressed let-7i-5p, the cell motility was decreased comparing to control pathogenic fibroblasts with no let-7i-5p change (Figure 3, lower right panel and lower left panel), while TGF-β1 and MMP2 protein levels were actually higher in let-7i-5p overexpressed pathogenic cells than that in control pathogenic cells (Figure 4A).

Interestingly, the protein expression pattern of TGF-βR1 mimics the results of motility analysis for corresponding cell types (Figure 3, 4), which indicates a potential role of TGF-beta receptor in regulation of cell motility. However, the protein expression of its ligand, TGF-beta1, is complimentary to that of TGF-beta receptor1 (Figure 6). This finding strongly suggests that the TGF-beta ligand expression is in a negative feedback loop to suppress the TGF-beta receptor level change and is consistent with the overall effect of compromising microRNA induced regulation at mRNA level and stabilizing the intracellular molecular status. In summary, let-7i-5p may regulate TGF-β1 and MMP2 as a group, and this regulation is hampered in let-7i-5p overexpressed pathogenic cells, while TGF-betaR1 was regulated by let-7i-5p in a separate pathway, and the regulation is independent of the pathogenic status of the cells.

### Let-7i-5p regulates TGF-beta signaling, fibroblast proliferation and cell morphology through Fibronectin, actin, TGFbetaR1 and TIMP1

The third group of targets regulated by let-7i-5p contains FN1, ACTIN, TGFBR1 and TIMP1, which are overexpressed at mRNA level upon dysregulation of let-7i-5p (no matter let-7i-5p’s expression is increased or decreased) (Figure 4A). The consistent pattern in normal and pathogenic fibroblasts implies that the fibrotic status of cells doesn’t affect the regulation by let-7i-5p. Given the fact that fibronectin and actin are essential for cell shape maintenance, it is not surprising that the regulatory effects by let-7i-5p at mRNA level was almost completely disappeared at protein level, strongly suggesting that the intracellular stabilization mechanism serves an important role to correct the abnormal gene expressions resulted from miRNA regulation during translation. Thus, if there is any regulation of fibronectin and actin, it is more likely to be spatial instead of quantitative, as their protein levels remain pretty stable with let-7i-5p level changes (Figure 4A). It would be interesting to check whether interrupted expression of let-7i-5p could induce microtubule and actin cytoskeleton reorganization.

The regulation of TGF-beta receptor1 at mRNA level was diminished at protein level, yet it still remains the same regulatory pattern as that in normal cells. However, in pathogenic cells, the effect is reversed, especially when overexpression of let-7i-5p resulted in a reduction of TGF-beta receptor 1 at protein level, which is completely opposite from the increase of TGFbetaR1 at mRNA level. This suggests that regulation of TGFbetaR1 by let-7i-5p might depend on fibrotic condition of the cells.

Manipulation of let-7i-5p expression causes little change of actin at protein level (Figure 4A), so if let-7i-5p functions through actin, it is more likely to be of actin distribution rather than of its protein quantity. It seems that normal let-7i-5p regulation is required for morphology control in normal fibroblasts (Figure2), yet the mechanism remains unclear. In literature, let-7i-5p dysregulation phenotype mimics that of Dematin mutations[17]. Dematin is an actin binding/bundling protein that regulates FAK activation through RhoA and regulate cell morphology[17] and is predicted to be a conserved target of miR181a-5p, miR181b-5p, miR181c-5p, miR181d-5p, and miR-4262 in human, yet little is known about its regulation by those miRNAs. In addition, it was reported that MMP1 overexpression could suppress Thioacetamide (TAA)-induced liver fibrosis in rat model[22]. TIMP1’s function in fibrosis has been in argument as the evidences are divergent from different studies, although many of the results suggests that its expression has no effect on fibrosis[23].

Interestingly, the protein levels of MMP1 and TIMP1 showed an increase with both up- and down-regulated let-7i-5p expression in normal cells, yet had little change in pathogenic cells, which is similar to the morphology changes in those cell types. This may imply a post-translational regulatory role of let-7i-5p in MMP1/TIMP1 protein expression and corresponding cell morphology. We cannot rule out the possibility that MMP1 and TIMP1 serve as downstream regulators of cell morphology in normal cells, and that such regulation is lost under fibrotic conditions.

### Regulation of let-7i-5p is partially dependent on pathogenic status of fibroblasts

In pathogenic fibroblasts, overexpression of let-7i-5p resulted in dramatic mRNA overexpression of all the molecular targets that were evaluated, and the differences are statistically significant (Figure 4A). This suggests that pathogenic fibroblasts might have increased sensitivity to let-7i-5p as a positive regulator at mRNA level. Similar regulation is not observed in the healthy/normal fibroblasts, which indicates that healthy cells could suppress the effects resulted from let-7i-5p overexpression, while pathogenic cells are unable to exert the suppression. This regulatory loss may associate with fibrosis or wound healing process in the pathogenic cells.

### Let-7i-5p and its function in post-translational regulation

Hsa-let-7i-5p is predicted to target on several post-translational pathways, such as protein deneddylation (through predicted target genes LRIG3, COPS6, GPS1, COPS8, etc.) and gene silencing by RNA/miRNA (through target genes AGO1, TRIM71, CELF1, etc.), and also in homeostatic process (through predicted target genes TLR4 and IGF1, members of HIF-1 signaling pathway[24]) (Figure 7). In this study, our data indicated an obviously compromised regulation at protein level by let-7i-5p expression manipulation comparing to that at mRNA level (Figure 5), suggesting a post-translational regulation and homeostatic maintenance. This may also reflect the partial co-efficiency of mRNA and protein expression, as well as the possibility that some mRNA may serve distinct functions without translated to protein, both on quantity and quality basis[25]. Further analysis needs to be performed to confirm whether let-7i-5p serves as a regulator during the process or not, and if yes, whether let-7i-5p regulate the translation in a feedback loop, or through an independent process.

### Potential clinical applications of hsa-let-7i-5p and microRNAs in PFUDD and urological diseases

miRNAs have been discussed as potential therapeutic targets and clinical biomarkers in various diseases [26-28]. microRNAs are under investigation in a number of recent clinical trials for various urologic complications (e.g. urinary bladder neck obstruction, urolithiasis, urinary tract disorders, renal carcinoma, kidney injury, etc.. (Table V), mainly by microRNA profiling in patients, with a few extended into studies targeting on a specific microRNA (such as miR-21). A recent clinical trial (NCT02639923) evaluates the correlation between serum let-7i expression and intracranial traumatic lesions, which is based on evidence from animal models[29].

**Table V.**
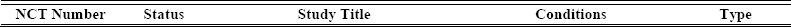
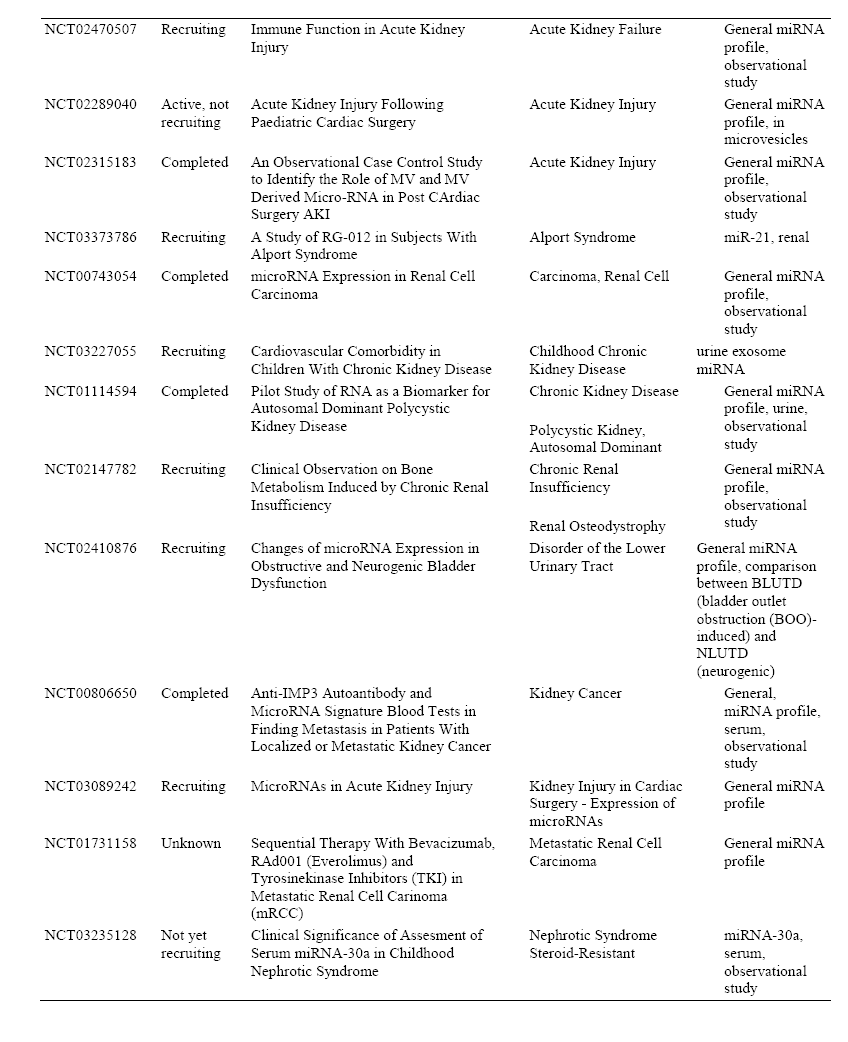
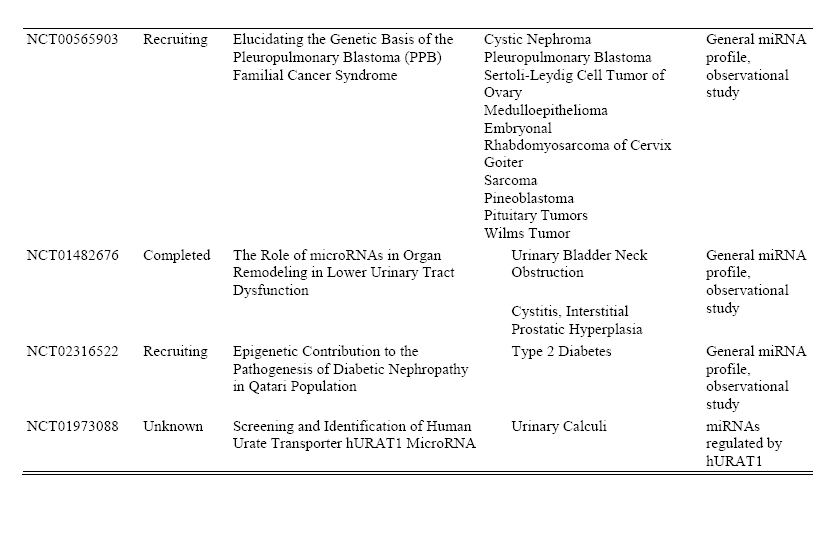
miRNA associated clinical trials in urological diseases

The unique regulatory functions of let-7i-5p in fibroblast proliferation, ECM regulation and homeostasis makes it an interesting drug target for complications involved with fibrosis, tissue reconstruction and cellular stress (Figure 7). We hope the data from this study could broaden our understanding of the function of hsa-let-7i-5p in normal and pathogenic fibroblasts and urethral tissues, so as to facilitate clinical diagnosis, treatment as well as tissue engineering as follow-up options for patients with PFUDD or other urological diseases.

## Conclusions

In this study, we analyzed let-7i-5p and its potential downstream targets (COL1A1, COL3A1, ELN, MMP1, VIM, FN1, ACTIN, TGFBR1, TIMP1, MMP2) in both normal and pathogenic fibroblasts. We found that let-7i-5p could regulate various signaling pathways and serve distinct functions in different cellular events, including tissue plasticity, cell motility, cell morphology, etc.. By comparing the regulatory effects at mRNA and protein levels, we extended our understanding of subcellular homeostasis and self-correction by stabilization mechanism during translation process. We conclude that let-7i-5p is a multi-functional regulator in cell signaling transduction, and it could be affected by fibrosis and pathogenic status.

## Acknowledgements

Financial support was obtained from the National Natural Science Fund of China (grant nos. 81470917 and 81700590), the Science and Technology Commission of Shanghai (grant no. 14JC1492100), and the Shanghai JiaoTong University Biomedical Engineering Cross Research Foundation (grant no. YG2014ZD07).

## References

1. Zhang K, Chen J, Zhang D, Wang L, Zhao W, Lin DY, Chen R, Xie H, Hu X, Fang X, Fu Q: microRNA expression profiles of scar and normal tissue from patients with posterior urethral stricture caused by pelvic fracture urethral distraction defects. Int J Mol Med 2018, 41:2733–2743.

2. Smith TG: Current management of urethral stricture disease. Indian Journal of Urology: IJU: Journal of the Urological Society of India 2016, 32:27–33.

3. Zhao J, Ren L, Liu M, Xi T, Zhang B, Yang K: Anti-fibrotic function of Cu-bearing stainless steel for reducing recurrence of urethral stricture after stent implantation. Journal of Biomedical Materials Research Part B: Applied Biomaterials:n/a–n/a.

4. O’Reilly S: MicroRNAs in fibrosis: opportunities and challenges. Arthritis Research & Therapy 2016, 18:11.

5. Jiang X-P, Ai W-B, Wan L-Y, Zhang Y-Q, Wu J-F: The roles of microRNA families in hepatic fibrosis. Cell & Bioscience 2017, 7:34.

6. Zhu H, Luo H, Zuo X: MicroRNAs: their involvement in fibrosis pathogenesis and use as diagnostic biomarkers in scleroderma. Experimental & Molecular Medicine 2013, 45:e41.

7. Vettori S, Gay S, Distler O: Role of MicroRNAs in Fibrosis. The Open Rheumatology Journal 2012, 6:130–139.

8. Jiang X, Tsitsiou E, Herrick SE, Lindsay MA: microRNAs and the regulation of fibrosis. The FEBS journal 2010, 277:2015–2021.

9. Rajasekaran S, Rajaguru P, Sudhakar Gandhi PS: MicroRNAs as potential targets for progressive pulmonary fibrosis. Frontiers in Pharmacology 2015, 6.

10. Bagnato G, Roberts WN, Roman J, Gangemi S: A systematic review of overlapping microRNA patterns in systemic sclerosis and idiopathic pulmonary fibrosis. European Respiratory Review 2017, 26.

11. Li H, Zhao X, Shan H, Liang H: MicroRNAs in idiopathic pulmonary fibrosis: involvement in pathogenesis and potential use in diagnosis and therapeutics. Acta Pharmaceutica Sinica B 2016, 6:531–539.

12. Li H, Zhao X, Shan H, Liang H: MicroRNAs in idiopathic pulmonary fibrosis: involvement in pathogenesis and potential use in diagnosis and therapeutics. Acta Pharmaceutica Sinica B 2016, 6:531–539.

13. Livak KJ, Schmittgen TD: Analysis of Relative Gene Expression Data Using Real-Time Quantitative PCR and the 2−ΔΔCT Method. Methods 2001, 25:402–408.

14. Reinhart BJ, Slack FJ, Basson M, Pasquinelli AE, Bettinger JC, Rougvie AE, Horvitz HR, Ruvkun G: The 21-nucleotide let-7 RNA regulates developmental timing in Caenorhabditis elegans. Nature 2000, 403:901.

15. O’Connell RM, Rao DS, Baltimore D: microRNA Regulation of Inflammatory Responses. Annual Review of Immunology 2012, 30:295–312.

16. Ludwig N, Leidinger P, Becker K, Backes C, Fehlmann T, Pallasch C, Rheinheimer S, Meder B, Stähler C, Meese E, Keller A: Distribution of miRNA expression across human tissues. Nucleic Acids Research 2016, 44:3865–3877.

17. Mohseni M, Chishti AH: The Headpiece Domain of Dematin Regulates Cell Shape, Motility, and Wound Healing by Modulating RhoA Activation. Molecular and Cellular Biology 2008, 28:4712–4718.

18. Vu LT, Keschrumrus V, Zhang X, Zhong JF, Su Q, Kabeer MH, Loudon WG, Li SC: Tissue Elasticity Regulated Tumor Gene Expression: Implication for Diagnostic Biomarkers of Primitive Neuroectodermal Tumor. PLoS ONE 2015, 10:e0120336.

19. Tao R, Fan X-X, Yu H-J, Ai G, Zhang H-Y, Kong H-Y, Song Q-Q, Huang Y, Huang J-Q, Ning Q: MicroRNA-29b-3p prevents Schistosoma japonicum-induced liver fibrosis by targeting COL1A1 and COL3A1. Journal of Cellular Biochemistry:n/a–n/a.

20. Wu F, Chakravarti S: Differential Expression of Inflammatory and Fibrogenic Genes and Their Regulation by NF-κB Inhibition in a Mouse Model of Chronic Colitis. The Journal of Immunology 2007, 179:6988.

21. Yang C, Zeisberg M, Mosterman B, Sudhakar A, Yerramalla U, Holthaus K, Xu L, Eng F, Afdhal N, Kalluri R: Liver fibrosis: Insights into migration of hepatic stellate cells in response to extracellular matrix and growth factors. Gastroenterology 2003, 124:147–159.

22. Iimuro Y, Nishio T, Morimoto T, Nitta T, Stefanovic B, Choi SK, Brenner DA, Yamaoka Y: Delivery of matrix metalloproteinase-1 attenuates established liver fibrosis in the rat. Gastroenterology 2003, 124:445–458.

23. Giannandrea M, Parks WC: Diverse functions of matrix metalloproteinases during fibrosis. Disease Models & Mechanisms 2014, 7:193–203.

24. Prabhakar NR, Semenza GL: Oxygen Sensing and Homeostasis. Physiology 2015, 30:340–348.

25. Cenik C, Cenik ES, Byeon GW, Grubert F, Candille SI, Spacek D, Alsallakh B, Tilgner H, Araya CL, Tang H, et al: Integrative analysis of RNA, translation, and protein levels reveals distinct regulatory variation across humans. Genome Research 2015, 25:1610–1621.

26. Christopher AF, Kaur RP, Kaur G, Kaur A, Gupta V, Bansal P: MicroRNA therapeutics: Discovering novel targets and developing specific therapy. Perspectives in Clinical Research 2016, 7:68–74.

27. Ji W, Sun B, Su C: Targeting MicroRNAs in Cancer Gene Therapy. Genes 2017, 8:21.

28. Mlcochova H, Hezova R, Meli AC, Slaby O: Urinary MicroRNAs as a New Class of Noninvasive Biomarkers in Oncology, Nephrology, and Cardiology. In RNA Interference: Challenges and Therapeutic Opportunities. edited by Sioud M. New York, NY: Springer New York; 2015: 439–463

29. Balakathiresan N, Bhomia M, Chandran R, Chavko M, McCarron RM, Maheshwari RK: MicroRNA Let-7i Is a Promising Serum Biomarker for Blast-Induced Traumatic Brain Injury. Journal of Neurotrauma 2012, 29:1379–1387.

